# Immunogenicity and protective efficacy of an influenza virus-like particle-based SARS-CoV-2 hybrid vaccine candidate in rhesus macaques

**DOI:** 10.1101/2024.05.24.595657

**Authors:** Sheikh Abdul Rahman, Ramireddy Bommireddy, Nanda Kishore Routhu, Lilin Lai, Christopher D. Pack, Sampath Ramachandiran, Mehul S. Suthar, Shaker J. C. Reddy, Periasamy Selvaraj, Rama Rao Amara

## Abstract

Severe acute respiratory syndrome coronavirus 2 (SARS-CoV-2) and influenza virus co-infections present a heightened COVID-19 disease and hospitalization cases. Here, we studied the immunogenicity and efficacy of an influenza-A/PR8 virus-like particle (^Flu^VLP)-based hybrid vaccine candidate displaying GPI-anchored SARS-CoV-2 receptor binding domain fused to GM-CSF and GPI-anchored interleukin-12 (^Flu^VLP-RBD) in rhesus macaques. Animals (n=4/group) received two doses of either ^Flu^VLP or ^Flu^VLP-RBD vaccine four weeks apart and were challenged with SARS-CoV-2 (WA1/2020) infection via intranasal and intratracheal routes. We determined vaccine-induced IgG and neutralizing antibody titers in serum and their association with viral replication in the lower and upper airways (lung, throat, and nose) and lung-associated pathologies. ^Flu^VLP-RBD vaccine induced a strong binding IgG in serum against multiple SARS-CoV-2 variants (WA1/2020, Delta and Omicron; BA.1). Both vaccines induced strong influenza A/PR8-specific IgG. Following the SARS-CoV-2 challenge, all four animals in the ^Flu^VLP-RBD group showed a profound control of virus replication in all three airway compartments as early as day 2 through day 10 (day of euthanasia). This level of viral control was not observed in the ^Flu^VLP group as 2-3 animals exhibited high virus replication in all three airway compartments. The protection in the ^Flu^VLP-RBD vaccinated group correlated positively with post challenge neutralizing antibody titer. These results demonstrated that a ^Flu^VLP-based hybrid SARS-CoV-2 vaccine induces strong antibody responses against influenza-A/PR8 and multiple SARS-CoV-2 RBD variants and protects from SARS-CoV-2 replication in multiple compartments in macaques. These findings provide important insights for developing multivalent vaccine strategies for respiratory viruses.

**Importance:** Co-infection with multiple respiratory viruses poses a greater risk than individual infections, especially for individuals with underlying health conditions. Studies in humans consistently demonstrated that simultaneous infection with SARS-CoV-2 and influenza leads to more severe respiratory illness and an increased rate of hospitalization. Therefore, developing hybrid vaccines targeting multiple respiratory viruses is of high importance. The hybrid vaccines also help to reduce the economic and logistic burden associated with vaccine coverage, distribution and storage. Here, we evaluate the immunogenicity and effectiveness of a novel hybrid flu-SARS-CoV-2 vaccine candidate using a nonhuman primate pre-clinical model. Our findings reveal that this vaccine elicits a strong immune response against influenza and SARS-CoV-2 viruses. Importantly, it provides strong protection against SARS-CoV-2 infection and associated pathological conditions.

## Introduction

The coronavirus disease of 2019 (COVID-19) remains a public health concern. Since the onset of the severe acute respiratory syndrome coronavirus 2 (SARS-CoV-2) pandemic, more than 700 million people have been infected, and nearly 7 million people have died worldwide. SARS-CoV-2 is an enveloped virus expressing spike (S) glycoprotein having receptor binding domain (RBD) region directly responsible for the virus attachment with the host receptor ACE2, fusion with the membrane, and entry into the cell causing infection (1, 2). Therefore, S or just the RBD represents a potent vaccine target, and indeed, targeting S/RBD as a vaccine antigen resulted in the development of multiple efficacious vaccines against SARS-CoV-2 quickly (3, 4). Several vaccine platforms have shown robust protection from severe illness and hospitalizations (5–10). However, the limited durability of vaccine-induced neutralizing antibody response and constant evolution of the SARS-CoV-2 virus has resulted constant emergence of variants of concern (VOC) necessitating repeated boosting of immune response either with wildtype spike (S) or S-variant vaccines. Moreover, co-infection with other respiratory viruses complicates clinical outcomes of SARS-CoV-2 infection. Therefore, developing novel vaccine strategies where the S protein can be easily modified to match the emerging VOCs and have the potential to target multiple respiratory viruses is of crucial importance.

SARS-CoV-2 comorbidities present a serious threat to the public health crisis (11, 12). In this regard, COVID-19 disease severity has been shown to aggravate under the CoV-2/flu co-infection condition, resulting in more significant hospitalizations and deaths (13, 14). Though limited, studies have shown that compared to individual SARS-CoV-2 or flu infection, co-infection results in greater infection persistence, inflammation, and lung damage (13). Given that ongoing SARS-CoV-2 infection overlaps with the flu season, co-immunization against SARS-CoV-2 and influenza might present a promising strategy to simultaneously protect from both respiratory infections. Here, limited studies have shown that either co-administration of SARS-CoV-2 and flu vaccines or a single vaccine co-expressing both SARS-CoV-2 and flu antigens induce strong immune response against both infections and provide protection against viral challenge in mice (15, 16). Consistently, co-administration of SARS-CoV-2 and flu vaccines in human ACE2 transgenic mice protected mice from SARS-CoV-2 and influenza virus challenge (17). Another study using a single vaccine based on live attenuated influenza A (LAIV) expressing SARS-CoV-2 RBD protected BALB/c mice from SARS-CoV-2 challenge (18). These studies pointed to the feasibility of developing a single vaccine platform for co-delivering more than one immunogen.

Vaccine platform plays an important role in the magnitude and quality of immune response induced. For example, mRNA vaccines induced greater antibody response and exhibited superior vaccine efficacy than the adenoviral vector vaccines (19, 20). Similarly, multiple factors can influence vaccine efficacy, such as delivery method, antigen presentation, and immune activation. Thus, immunogen delivery vehicles, immunogen design, and type of adjuvants significantly affect the magnitude and durability of immune responses. Biological adjuvants co-expressed with the immunogens improve vaccine effects (2, 21–25). These adjuvants potentiate the Th1 immune response, ensuring robust anti-viral immunity. Consistently, IL-12 has been shown to enhance clinical benefit by promoting Th1 response in cancer patients (22, 26–28). Similarly, GM-CSF has been shown to improve immune response primarily by potentiating dendritic cells (29, 30). Our previous studies have also demonstrated that an influenza-virus-like particle-based vaccine co-displaying GPI-RBD-GM-CSF fusion protein and GPI-IL-12 protected mice from mouse-adapted SARS-CoV-2 and influenza (31). Here, we report the immunogenicity and efficacy of this novel influenza-VLP-based SARS-CoV-2 dual vaccine candidate in the nonhuman primate model.

## Results

### Influenza-A/PR8 virus-like particle (VLP)-incorporated SARS-CoV-2 receptor binding domain (RBD) retains functional activity

Influenza-A/PR8 virus-like particle (VLP)-based SARS-CoV-2 receptor binding domain (RBD) hybrid vaccine (Figure 1A) was constructed and characterized as shown previously (31). Receptor binding activity of the recombinant RBD was confirmed by ACE2-binding ELISA and MM57 mAb binding ELISA (Figure 1B). The flow cytometry analysis confirmed the successful incorporation of the GPI-RBD-hGM-CSF and GPI-IL-12 into VLPs (Figure 1E, D).

**Figure 1.**
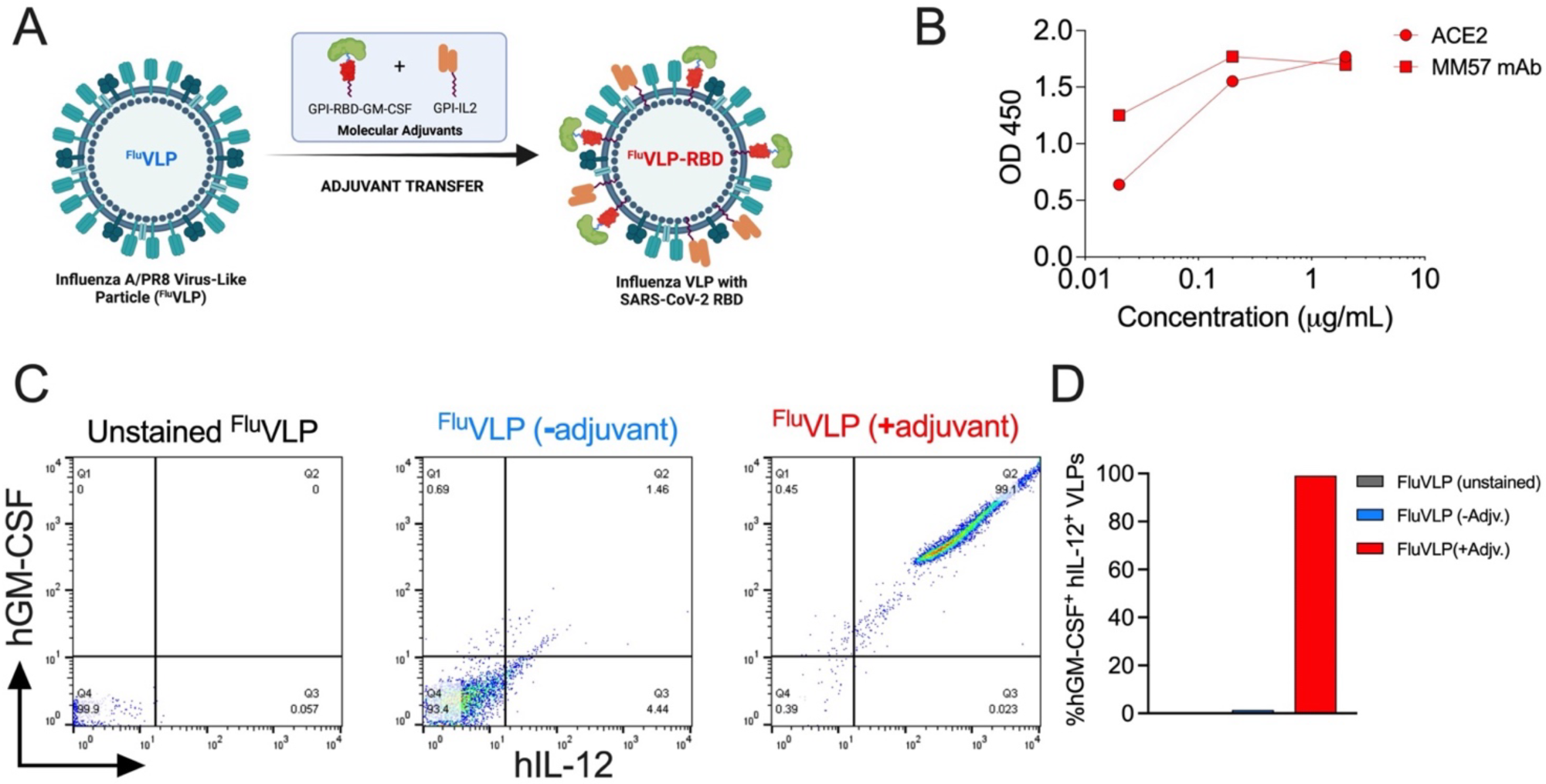
Vaccine design and characterization of fusion protein (A-C). (**A**) Design of Influenza A/PR8 virus like protein expressing SARS-CoV-2 receptor binding domain and indicated molecular adjuvants. (**B**) Validation of ACE2-binding ability of fusion GPI-RBD-hGM-CSF protein by ELISA. (**C**) Validation of protein transferred GPI-RBD-GM-hCSF and hIL-12 incorporated VLPs using flow cytometry. (**D**).

To test the immunogenicity and the protective efficacy of ^Flu^VLP-RBD hybrid vaccine in a preclinical nonhuman primate model, we employed eight Indian-origin rhesus macaques and divided them equally into two vaccine groups (**Figure 2A**). One group was immunized with 25 μg of unmodified ^Flu^VLP, whereas the second group was immunized with ^Flu^VLP incorporated with GPI-RBD-GM-CSF plus GPI-IL-12 (^Flu^VLP-RBD hybrid vaccine). The booster doses were given four weeks post prime doses. Blood was collected at week 0 and 2 of each vaccine dose. Both vaccinated groups were challenged with 10^5^ pfu SARS-CoV-2 Washington-1/2020 strain via intra-nasal and intra-tracheal routes at week eight of first vaccine dose. The protective efficacy of the vaccines was assessed by measuring sub-genomic SARS-CoV-2 RNA in bronchioalveolar lavage, nasopharynx, and trachea at day 0, 2, 4, 7, and 10 of viral challenge. Histopathology was analyzed from lung tissues at the necropsy. We compared viral replication and histopathology with 7 historical control rhesus macaques from our recently published reports (32, 33). Out of 7 historical control group animals, 5 macaques received wild type modified vaccinia Ankara (MVA) vaccine and 2 macaques were left untreated.

**Figure 2.**
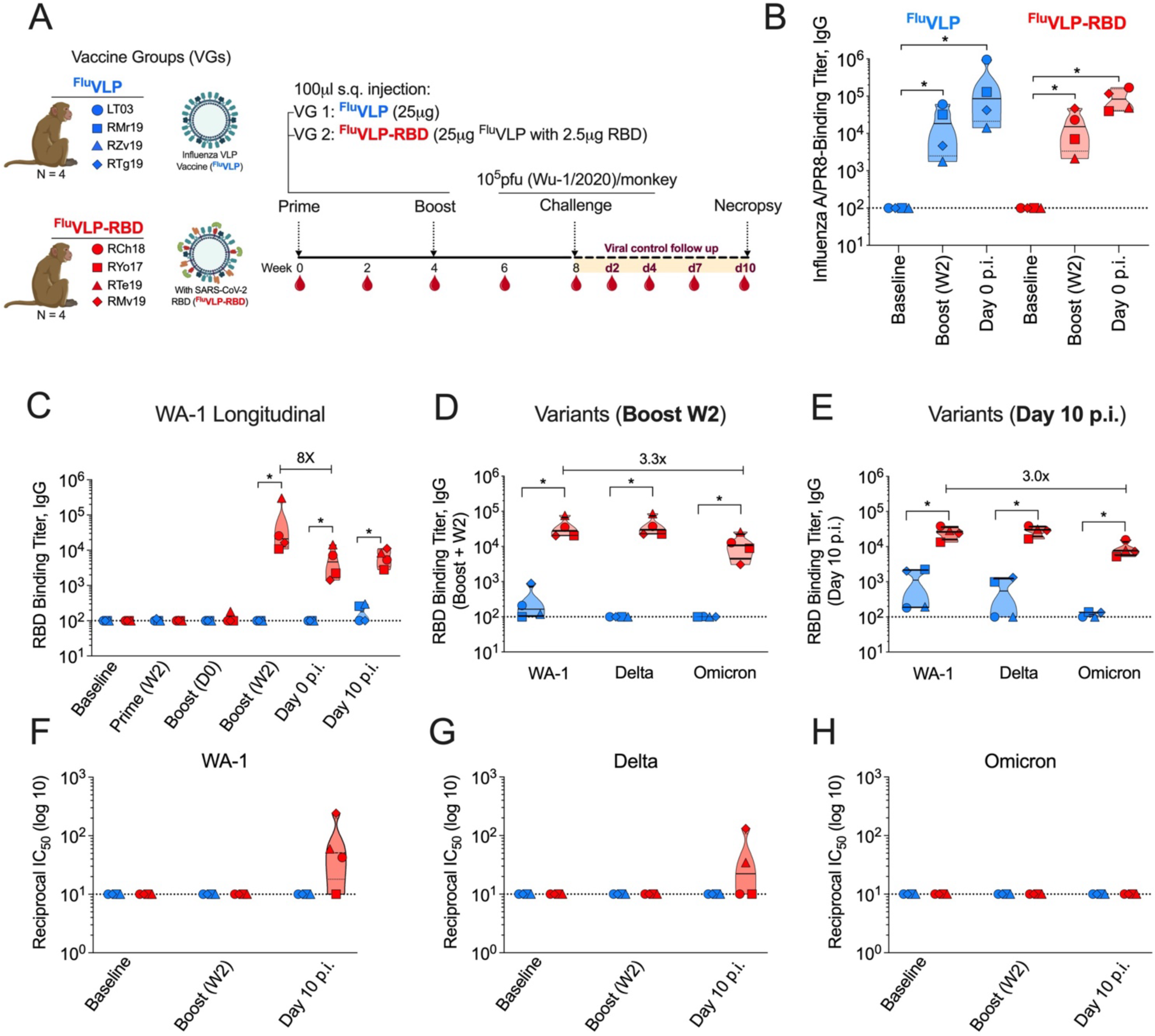
NHP study design and antibody responses elicited by influenza A/PR8 VLP-SARS-CoV-2 RBD dual vaccine candidate (A-H). (**A**) Study design showing NHP treatment groups, immunogen design, vaccination regimen, blood collection and SARS-CoV-2 WA-1/2020 challenge scheme. (**B**) Influenza A/PR8-binding IgG titers at indicated timepoints post vaccination in ^Flu^VLP (in blue) and ^Flu^VLP-RBD (in red) groups. (**C**) Longitudinal SARS-CoV-2 WA-1/2020 strain RBD binding IgG titers shown for indicated timepoints in both vaccinated groups. (**D, E**) Binding IgG titers shown for SARS-CoV-2 variant Delta and Omicron compared to the ancestral WA-1 strain at day 0 of infection (**D**) and day 10 post infection (**E**). (**F-H**) Neutralizing antibody titer against WA-1/2020 strain is shown for WA-1 (**F**), Delta (**G**) and Omicron (**H**) strains. Statistical significance was calculated by Mann-Whitney U test. *P < 0.05.

### ^Flu^VLP with or without SARS-CoV-2 RBD induced robust IgG against influenza-A/PR8

To determine humoral response to influenza VLP induced by ^Flu^VLP compared with the ^Flu^VLP-RBD hybrid vaccine, we analyzed total immunoglobulin G (IgG) response generated against influenza A/PR8 VLP in sera obtained from vaccinated rhesus macaques at week 0, week 6 (boost + w2), and week 8 (day of SARS-CoV-2 challenge) of vaccinations (Figure 2A). ^Flu^VLP (with or without RBD) vaccinations induced significantly higher (∼113-fold) influenza-binding IgG compared with the baselines (**Figure 2B**). The magnitude of IgG induction was comparable between ^Flu^VLP (geomean 11305) and ^Flu^VLP-RBD hybrid vaccine (geomean 11,311) vaccinated macaques. Encouragingly, anti-influenza IgG remained persistent in ^Flu^VLP (geomean 92,844; 8-fold high compared to week 2 post boost) and ^Flu^VLP-RBD hybrid vaccine (geomean 78,818; 7-fold high compared to week 2 post boost) groups. These data showed a robust generation of anti-influenza binding antibodies, indicating no interference by SARS-CoV-2 RBD.

### ^Flu^VLP-RBD hybrid vaccine induced robust humoral response against multiple SARS-CoV-2 variants

We determined whether ^Flu^VLP-RBD hybrid vaccine-induced antibody binds to SARS-CoV-2 washington-1/2020, Delta, and Omicron variants by analyzing total IgG binding. In this direction, we tested sera from vaccinated groups (with or without RBD) at weeks 0, 2, 4, 6, 8 (day 0 of infection) and 9 (day 10 post infection, p.i.) post-vaccination. The ^Flu^VLP-RBD vaccine induced significantly higher (∼343-fold) RBD-binding IgGs (34324; geomean) compared with the ^Flu^VLP (**Figure 2C**). On the day of the SARS-CoV-2 challenge, the magnitude of RBD-binding IgG was consistently higher in ^Flu^VLP-RBD immunized macaques (4245; geomean) compared with the ^Flu^VLP immunized macaques (p = 0.02). We observed an 8-fold contraction in RBD-binding IgG on the day of challenge compared with the peak response at week 2 post boost. Encouragingly, RBD-binding IgG persistent over time until day 10 p.i. We observed low level induction of RBD-binding IgG (166; geomean) in ^Flu^VLP vaccinated macaques 10 days p.i. that was significantly lower (37-fold; p = 0.02) compared with the RBD-binding IgG observed in ^Flu^VLP-RBD vaccinated macaques (6130; geomean).

Next, to determine the breadth of ^Flu^VLP-RBD hybrid vaccine-induced antibody responses, we analyzed antibody binding against Delta and Omicron (BA.1) SARS-CoV-2 variants. At peak response (week 2) post boost, we observed robust induction of binding-IgGs against all indicated variants, significantly higher than the ^Flu^VLP vaccine immunized group (**Figure 2D, E**). At week 2 post boost, the magnitude of binding-IgG was comparable between WA-1/2020 (32489; geomean) and Delta strain (∼35660; geomean). However, the magnitude of RBD-binding-IgG to Omicron (BA.1) was ∼3-fold less compared with the WA-1/2020 (9717; geomean). At day 10 post infection the magnitude of binding-IgG was comparable between WA-1/2020 (∼24345; geomean) and Delta strain (∼27698; geomean). However, the magnitude of RBD-binding-IgG to Omicron (BA.1) was ∼3-fold less compared with the WA-1/2020 (∼8314; geomean) (**Figure 2E**). The IgG response on day 10 post challenge was comparable with the response on the day of the challenge against all indicated variants. These data demonstrate a robust induction of humoral response to multiple variants of the SARS-CoV-2 RBD, indicating a broad IgG response by ^Flu^VLP-RBD hybrid vaccine.

### SARS-CoV-2 WA-1/2020 challenge induced neutralizing antibody in ^Flu^VLP-RBD vaccinated rhesus macaques

We next evaluated virus neutralizing potential of the vaccine-induced humoral responses. We tested serum collected at week 0 (baseline), 6 (boost + w2), and 9.3 (day 10p.i.) of vaccinations and determined neutralizing antibody (nAb) using mNeonGreen SARS-CoV-2 virus and defined 50% reduction in focus reduction neutralization titer (FRNT) (Figure 2F-H). We observed no nAb in sera from macaques at peek timepoint post vaccination (**Figure 2F**). Encouragingly, upon viral challenge, only ^Flu^VLP-RBD hybrid vaccinated macaques showed nAb in 3 out of 4 macaques ranging from 10-243 (geomean ∼50) but not the ^Flu^VLP group (**Figure 2F**). We further evaluated the neutralizing antibody response against the Delta variant. Similar to the response observed against WA-1/2020 strain, no neutralization was detected after vaccination, but 2 out of 4 macaques showed nAb induction only upon virus challenge (**Figure 2G**). We also analyzed nAb against the Omicron variant. However, there was no response observed (**Figure 2H**). These data indicated that though immunization with ^Flu^VLP-RBD hybrid vaccine failed to induce measurable neutralizing antibody, it has the ability to generate nAb post viral challenge. Moreover, the nAb had potential to cross-neutralize Delta variant but not Omicron.

### ^Flu^VLP-RBD hybrid vaccine did not induce robust CD4 and CD8 T cell responses

In order to determine T cell induction potential of the ^Flu^VLP-RBD hybrid vaccine platform we analyzed cytokine expression in antigen specific CD4 and CD8 T cell subsets. We analyzed combined response to spike RBD domain peptide pools in PBMCs at week 0 (baseline), 2 (prime + week 2), 6 (boost + week 2), and 9.3 (day 10p.i.) of immunization. We did not observe robust induction of either IFN γ+, IL2+, or TNFα+ expressing CD4 T cells post prime/boost immunization, however both groups induced antigen-specific cytokine (IFN γ+, IL2+, and TNFα+) responses upon viral challenge (**Figure 3A, B**). Post challenge ^Flu^VLP vaccinated group showed higher CD4 T cell IFNγ+ (0.08; geomean), IL-2 (0.07; geomean), and TNFα+ (0.05; geomean) responses compared with the IFNγ+ (0.03; geomean), IL-2 (0.035; geomean), TNFα+ (0.015; geomean) in ^Flu^VLP-RBD hybrid vaccine group. Though these responses were comparable between the groups, only CD4+IFNγ+ response in ^Flu^VLP-RBD group showed statistical significance compared with the baseline (**Figure 3B**). Next, we analyzed the polyfunctionality of the infection induced CD4 T cell responses in vaccinated groups. The ^Flu^VLP group showed CD4 T cells expressing two (IFNγ+TNFα+/ IFNγ+IL-2+/ TNFα+IL-2+) or all three (IFNγ+IL-2+TNFα+) cytokines highlighting their polyfunctional nature (**Figure 3C; blue**). Similar polyfunctionality was observed in ^Flu^VLP-RBD group, however the dual cytokine producers were markedly higher in ^Flu^VLP-RBD group (**Figure 3C; red**). We further analyzed antigen specific CD8 T cell cytokine response in blood. Interestingly, CD8 T cells in ^Flu^VLP group showed significantly higher (p = 0.028) IFN γ cytokine expression (0.08; geomean) in response to the S peptides compared with the ^Flu^VLP-RBD group (0.02; geomean) (**Figure 3D, E**). Both, IL-2 and TNFα expression was higher in ^Flu^VLP group compared with the ^Flu^VLP-RBD group. Finally, we analyzed polyfunctionality of the CD8 T cell responses. CD8 T cells in ^Flu^VLP group showed some degree of polyfunctionality as indicated by expression of only dual cytokine combinations but not all three cytokines together (**Figure, 3F; blue**). As indicated above ^Flu^VLP-RBD group showed low levels of only IFN γ+ expression but no other cytokines and thereby no polyfunctionality was observed (**Figure 3 F; red**). Taken together ^Flu^VLP-RBD hybrid vaccine failed to induce potent CD4 and CD8 T cell responses although CD4 and CD8 T cell responses were observed upon viral challenge in both vaccine groups. Quality of CD4 T cell responses was superior in ^Flu^VLP-RBD vaccine group over ^Flu^VLP group. Conversely, CD8 T cell response was superior in ^Flu^VLP vaccine group post infection compared with the ^Flu^VLP-RBD group.

**Figure 3.**
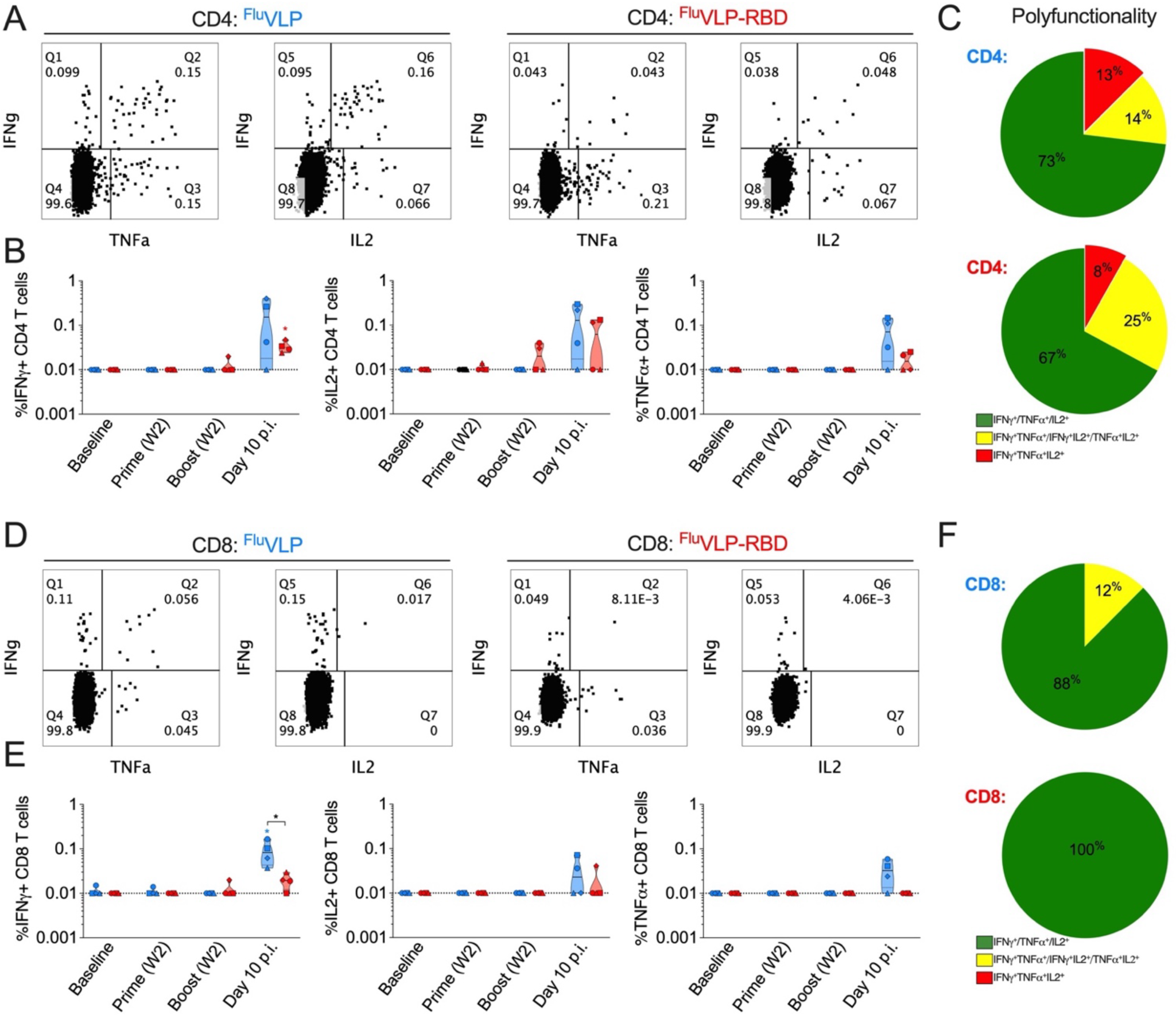
Spike specific T cell response in blood post vaccination and WA-1/2020 viral challenge (A-E). (**A**) Representative flow plots showing CD4 cytokine (IFN γ^+^, TNFα^+^, and IL2^+^) response to spike peptides in peripheral blood mononucleocytes (PBMCs) of ^Flu^VLP (in **blue**) and ^Flu^VLP-RBD (in **red**) vaccinated rhesus macaques at day 10 post infection (p.i.) **(B)** Longitudinal cytokine responses plot shown for IFN γ^+^, TNFα^+^, and IL2^+^ CD4 T cell responses at indicated time points. (**C**) Pie plot showing proportion of polyfunctional CD4 T cell response to S-peptides at day 10 p.i. as indicated by single (IFN γ^+^, TNFα^+^, and IL2^+^) cytokine response in green, double (IFN γ^+^TNFα^+^, IFN γ^+^IL2^+^and TNFα^+^IL2^+^) cytokine response in yellow, and triple (IFN γ^+^TNFα^+^IL2^+^) cytokine response in red. (**D**) Representative flow plots showing CD8 cytokine (IFN γ^+^, TNFα^+^, and IL2^+^) response to spike peptides in PBMCs of ^Flu^VLP (in **blue**) and ^Flu^VLP-RBD (in **red**) vaccinated rhesus macaques at day 10 p.i. **(E)** Longitudinal cytokine responses plot shown for IFN γ^+^, TNFα^+^, and IL2^+^ CD8 T cell responses at indicated time points. (**F**) Pie plot showing proportion of polyfunctional CD4 T cell response to S-peptides at day 10 p.i. as indicated by single (IFN γ^+^, TNFα^+^, and IL2^+^) cytokine response in green, double (IFN γ^+^TNFα^+^, IFN γ^+^IL2^+^and TNFα^+^IL2^+^) cytokine response in yellow, and triple (IFN γ^+^TNFα^+^IL2^+^) cytokine response in red. Statistical significance was calculated by Mann-Whitney U test. *P < 0.05.

### ^Flu^VLP-RBD hybrid vaccine protected macaques from SARS-CoV-2 challenge and severe viral replication in lungs, nasopharynx, and throat

We next analyzed the protective ability of the ^Flu^VLP-RBD hybrid vaccine platform against ancestral SARS-CoV-2 Washington strain (WA-1/2020). Here, rhesus macaques in both vaccinated groups (with and without RBD) were challenged at week 8 post vaccinations and samples from BAL, nasopharynx, and throat were analyzed for evidence of viral replication by analyzing sub-genomic viral RNA at day 0, 2, 4, 7 and 10 of viral challenge. The magnitude of viral replication was compared between the vaccinated groups and with a previously published ancestral control group (32, 33). ^Flu^VLP-RBD group showed robust protection from WA-1/2020 replication in all three compartments compared with other groups (**Figure 4A-I**). In BAL, ^Flu^VLP-RBD showed robust control of the viral replication (geomean: 17 copies/mL; range: undetectable – 949 copies/mL). Out of four macaques in ^Flu^VLP-RBD group two showed complete protection, one macaque showed lower magnitude of replication at day 2 only (949 copies/mL) and another macaque showed low grade blip at day 2 (136 copies/mL) and 7 (178 copies/mL) as opposed to the 6 out of 7 macaques in historical ancestral control showing high magnitude of viral replication (geomean: 162 copies/mL; range: undetectable – 629716 copies/mL). ^Flu^VLP vaccinated macaques also did not demonstrate robust control (geomean 141 copies/mL; range: undetectable – 820,000 copies/mL). Surprisingly, 2 out of 4 macaques in ^Flu^VLP group showed no virus replication (**Figure 4A, B**). Overall, magnitude of viral replication as indicated by area under curve showed 169-fold lower viral load in ^Flu^VLP-RBD compared with the unvaccinated ancestral control and 39-fold lower than the ^Flu^VLP group (**Figure 4C**).

**Figure 4.**
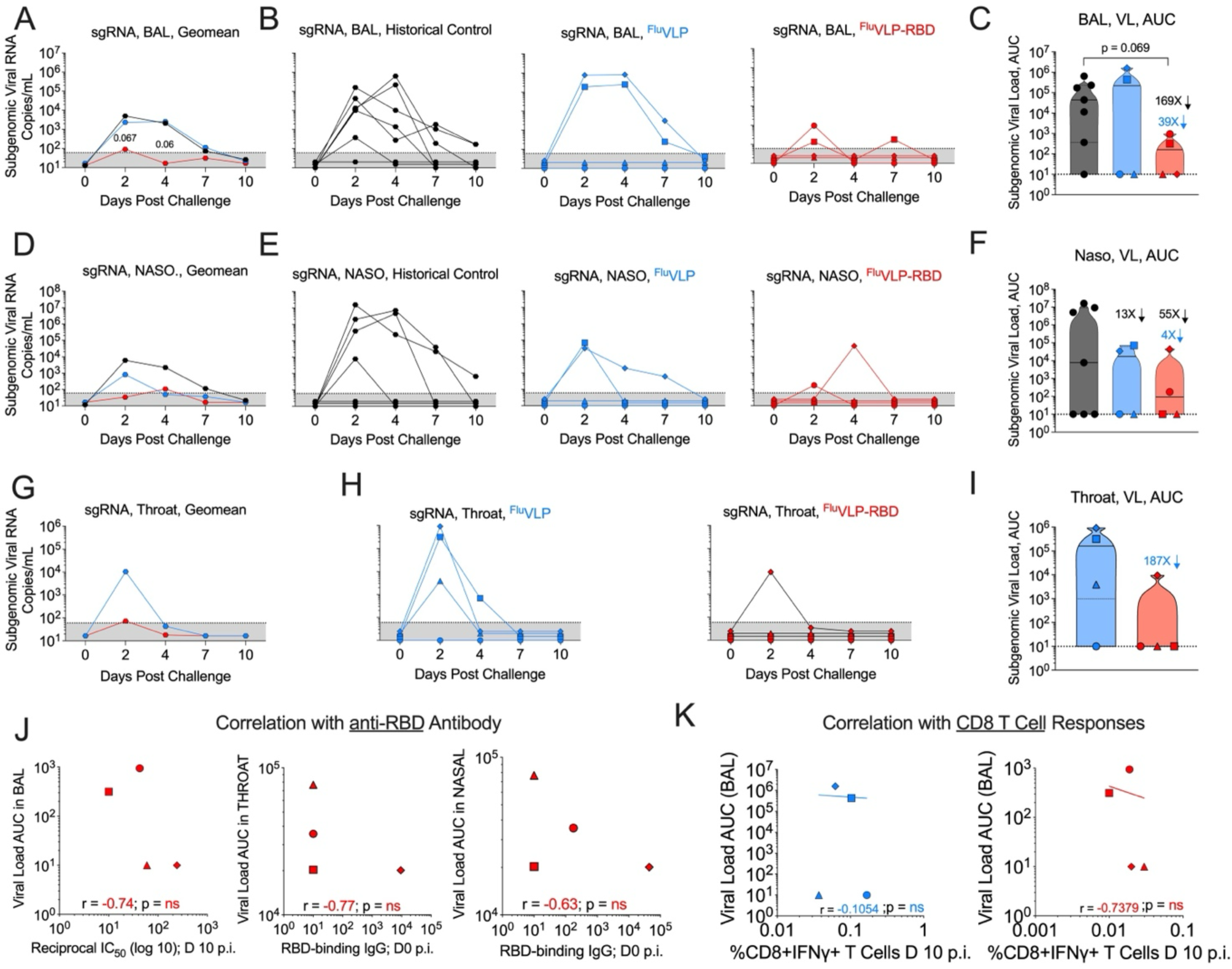
Protection against SARS-CoV-2 WA-1/2020 viral challenge and correlates of protection in rhesus macaques (A-K). (**A-C**) Longitudinal magnitude of viral load in buccopharyngeal lavage (BAL) followed for 10 days post infection (p.i.) represented as geomean (**A**), individual replication kinetics in historical unvaccinated **<u>historical control</u>** (black)^(32, 33)^, ^Flu^VLP (blue) and ^Flu^VLP-RBD (red) vaccinated macaques (**B**), and viral load area under curve from day 0 to 10 p.i. shown for each indicated groups (**C**). (**D-F**) Longitudinal magnitude of viral load in nasopharynx followed for 10 days p.i. represented as geomean (**D**), individual replication kinetics in unvaccinated **<u>historical control</u>** (black)^(32, 33)^ ^Flu^VLP (blue) and ^Flu^VLP-RBD (red) vaccinated macaques (**E**), and viral load area under curve from day 0 to 10 p.i. shown for each indicated groups (**F**). (**G-I**) Longitudinal magnitude of viral load in throat followed for 10 days p.i. represented as geomean (**G**), individual replication kinetics in ^Flu^VLP (blue) and ^Flu^VLP-RBD (red) vaccinated macaques (**H**), and viral load area under curve from day 0 to 10 p.i. shown for each indicated groups (**I**). (**J**) Correlation between neutralizing and RBD-binding antibodies with viral load magnitude in BAL, throat, and nasopharynx are shown for ^Flu^VLP-RBD vaccinated rhesus macaques. (**K**) Correlation between viral load area under curve in BAL with CD4 IFNγ response in blood at day 10 p.i. Statistical significance was calculated by Mann-Whitney U test. Correlation was determined using non-parametric Spearman correlation.

Next, we analyzed viral replication in the nasopharynx. In nasopharynx, ^Flu^VLP-RBD showed robust control of the viral replication (geomean: 18 copies/mL; range: undetectable – 44600 copies/mL). Out of four macaques in ^Flu^VLP-RBD group, two showed complete protection, one macaque showed a very low magnitude of replication at day 2 only (177 copies/mL) and another macaque showed replication at day 4 only (44600 copies/mL) as opposed to the 4 out of 7 macaques in historical ancestral control showing high magnitude of viral replication (geomean: 177; range: undetectable – 15740453 copies/mL). Interestingly, in ^Flu^VLP vaccinated group 2 out of 4 macaques showed no virus replication and 1 showed high and persistent viremia, whereas 1 showed moderate replication at day 2. Overall, magnitude of viral replication as indicated by area under curve showed 55-fold lower viral replication in ^Flu^VLP-RBD compared with the unvaccinated historical ancestral control and 4-fold lower compared with the ^Flu^VLP group (**Figure 4F**). We also analyzed viral replication in the throat. In ^Flu^VLP-RBD group 3 out of 4 macaques showed a complete absence of measurable viral load and only 1 macaque showed moderate viremia at day 2 only (9390 copies/mL) compared with the ^Flu^VLP group wherein 3 out 4 macaques showed a relatively higher degree of replication at day 2 (3900 copies/mL – 919000) (**Figure 4G, H**). Overall, ^Flu^VLP-RBD showed 187 times lower geomean viral load compared with the ^Flu^VLP group (**Figure 4I**). Encouragingly, the ^Flu^VLP-RBD group showed an inverse correlation between viral load and neutralizing that was induced post challenge as well as RBD-binding IgGs induced by ^Flu^VLP-RBD vaccination (**Figure 4J**). Encouragingly, viral control in ^Flu^VLP-RBD group showed inverse correlation with the antigen-specific CD8 T cells but not in ^Flu^VLP group (**Figure 4K**). Overall, these data showed robust protection from WA1/2020 in lungs, nose, and throat, and protection correlated with the antibody responses, though a statistical significance was not reached due to a small number of animals.

### ^Flu^VLP-RBD vaccinated macaques showed lower lung pathology and correlated with lower inflammatory immune response

To understand ^Flu^VLP-RBD mediated protection against lung pathology induced by SARS-CoV-2 replication, we analyzed multiple lung lobes for type 2 pneumocyte hyperplasia, alveolar septal thickening, and perivascular cuffing. A combined score was compared for the severity of lung pathology between both vaccinated groups and compared with the pathology observed in unvaccinated ancestral control macaques (**Figure 5A**). A scale between 0-5 (mild), 5-10 (moderate), and >15 (severe) was used to estimate the severity of the lung pathology. Consistent with the blunted viral replication ^Flu^VLP-RBD vaccinated macaques showed mild pathology (geomean 2; range 1-6), which was considerably lower (p = 0.06) compared with the pathology observed in historical ancestral control macaques (geomean 6; range 2 – 10) albeit statistically not significant due to small sample size. Pathology observed in ^Flu^VLP-RBD vaccinated macaques was also lower compared with the pathology observed in ^Flu^VLP immunized macaques (geomean 5; range: 1 – 15). Consistently, pathological conditions such as pneumocyte hyperplasia directly correlated with the viral replication in BAL of ^Flu^VLP-vaccinated macaques but not in ^Flu^VLP-RBD-vaccinated macaques (**Figure 5B**). Similarly, we observed direct correlation of CD4 T cell response post infection with the lung pathology only in ^Flu^VLP group (**Figure 5C**). Interestingly, CD8 T cell response post-infection directly correlated with the neutrophils before the challenge (**Figure 5D**), indicating infection and inflammation-induced cellular responses to SARS-CoV-2 replication in the ^Flu^VLP group. Again, the correlation was only observed in ^Flu^VLP and not in ^Flu^VLP-RBD vaccinated group. The histopathological condition directly correlated with the magnitude of viral load in the FluVLP group but not in the ^Flu^VLP-RBD group (**Figure 5E**). These findings showed that ^Flu^VLP-RBD hybrid vaccine protected macaques from severe lung pathology caused by SARS-CoV-2 infection.

**Figure 5.**
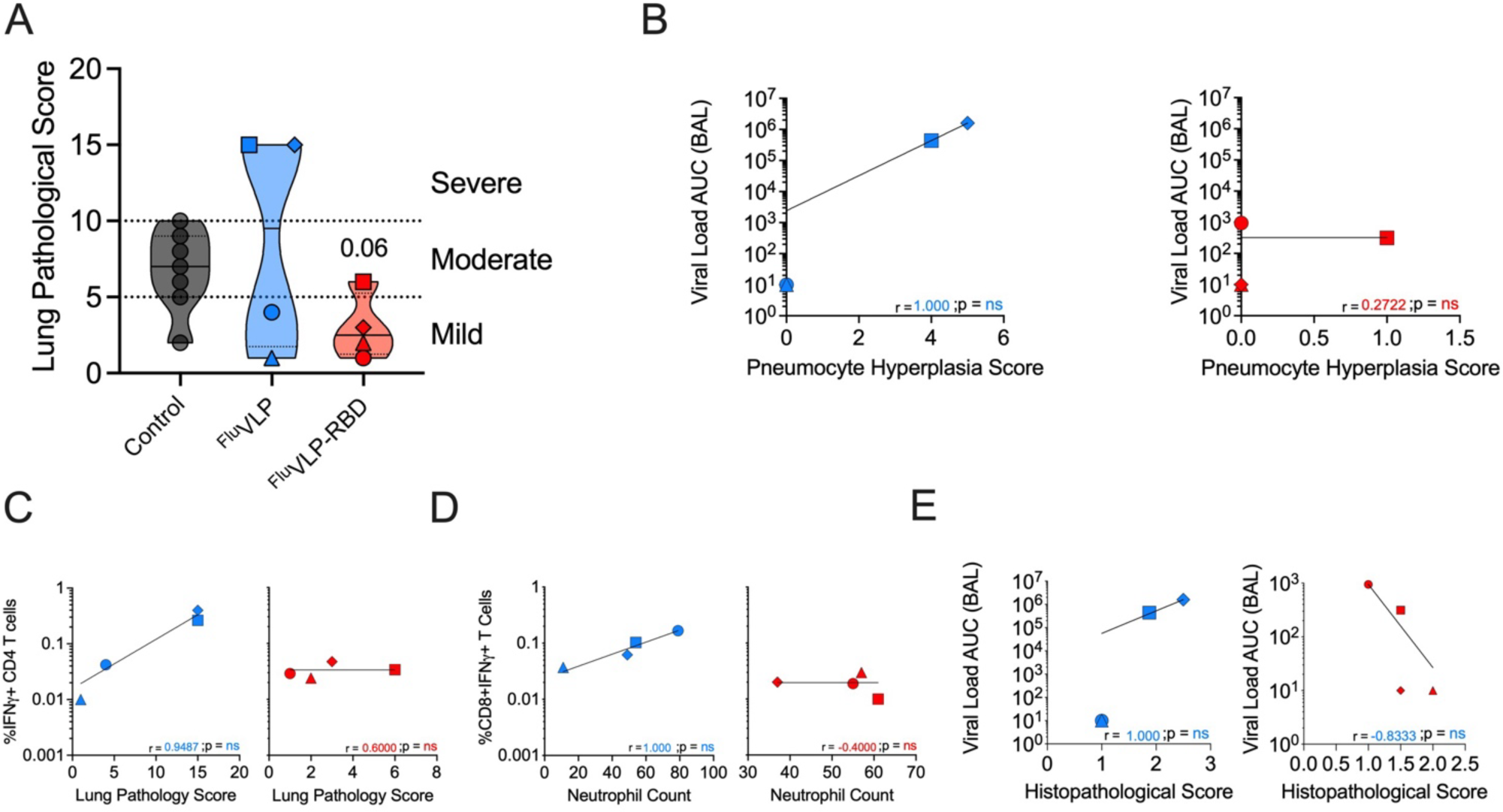
Histopathological condition and correlates of lung pathology post SARS-CoV-2 WA-1/2020 infection (A-D). (**A**) Magnitude of histopathological condition in lungs at day 10 post-infection (p.i.) is shown between **<u>historical control</u>** (**in black**)^(32, 33)^, ^Flu^VLP (**in blue**) and ^Flu^VLP-RBD (**in red**). (**B**) Correlation of lung pneumocyte hyperplasia with magnitude of virus replication in ^Flu^VLP (**in blue**) and ^Flu^VLP-RBD (**in red**) is shown. (**C**) Correlation of antigen specific CD4^+^IFNγ^+^ cytokine response and lung pathology in ^Flu^VLP (**in blue**) and ^Flu^VLP-RBD (**in red**) is shown. (**D**) Correlation of antigen-specific CD8^+^IFNγ^+^ cytokine response with magnitude of neutrophils prior to viral challenge (−2-week pre-challenge). (**E**) Correlation between viral load area under curve in BAL with histopathological score in ^Flu^VLP (**in blue**) and ^Flu^VLP-RBD (**in red**) vaccinated rhesus macaques. Correlation was determined using non-parametric Spearman correlation.

## Discussion

SARS-CoV-2 and influenza co-infection is a global concern because both viruses can cause respiratory infection and may lead to a more severe form of pneumonia, respiratory failure, and even death (11–14). A co-infection can be particularly detrimental for vulnerable populations such as the elderly, those with underlying conditions, and pregnant women. The risk of co-infection particularly aggravates during flu season. Therefore, there is a growing interest to co-immunize against both influenza and SARSCoV-2, either by co-delivering vaccines against both viruses individually and/or developing a dual vaccine for both viruses (16, 18, 31). To date, there is no preclinical data either in humans or nonhuman primates to support whether a dual vaccine co-expressing influenza and SARS-CoV-2 antigens will be safe and immunogenic and whether it will be protective. Here, we tested the immunogenicity and protective efficacy of a hybrid vaccine based on influenza-A/PR8-virus-like particle (^Flu^VLP) displaying SARS-CoV-2 RBD (^Flu^VLP-RBD) in rhesus macaques. Our results showed that the ^Flu^VLP-RBD hybrid vaccine induced a strong antibody response against influenza-A/PR8 and SARS-CoV-2. ^Flu^VLP without RBD showed comparable induction of binding IgG against influenza-A/PR8-VLP. However, we could not test the protective ability against influenza in the macaque model. Therefore, a qualitative difference cannot be ruled out and requires further validation.

The ^Flu^VLP-RBD hybrid vaccine induced strong antibody responses against multiple SARS-CoV-2 variants (WA-1/2020, Delta, and Omicron), indicating no interference of immunogen co-presentation on induction of either anti-influenza or anti-SARS-CoV-2 RBD humoral responses. We then showed that the ^Flu^VLP-RBD hybrid vaccine, despite not showing measurable neutralizing antibody response during vaccination, showed induction of neutralizing antibody upon SARS-CoV-2 challenge, and the post infection titer of neutralizing antibody correlated with the protection. It is noteworthy that a high magnitude of neutralizing antibody at peak vaccine response was not required and that an efficient priming might be sufficient for protection. Though this is an intriguing finding, more studies are needed to understand the detailed nature of immunological parameters modulated by the ^Flu^VLP-RBD hybrid vaccine carrying RBD immunogen and the underlying relation with the protection. We further showed that ^Flu^VLP-RBD did not induce CD4 as well as CD8 T cell responses. We suspect low levels of RBD alone might not be sufficient to induce a good cellular response. Though post viral challenge, both vaccinated groups showed comparable CD4 T cell cytokine responses. However, the ^Flu^VLP group showed significantly higher CD8+IFNγ+ response, which correlated with the magnitude of viral load. This correlation was not observed in the ^Flu^VLP-RBD hybrid vaccine indicating a higher infection level in the ^Flu^VLP group driving T cell responses. Further studies are needed to evaluate the T cell induction potential of the dual vaccine carrying higher RBD doses.

Interestingly, we observed partial protection against SARS-CoV-2 in macaques who received ^Flu^VLP vaccine that did not contain SARS-CoV-2 RBD. Surprisingly, protection directly correlated with influenza-A/PR8-specific binding IgG in these animals. Whether this is a direct effect or an indirect effect is yet to be determined. Emerging studies point to several reasons that might have contributed to the protection. Several studies have shown cross-protection from innate immunity against SARS-CoV-2 infection and severe clinical symptoms (34–45). However, precise mechanisms of influenza-SARS-CoV-2 cross-protection are yet to be defined. Limited studies have pointed to cross-reactive antibodies and CD8 T cells to SARS-CoV-2 and influenza (46, 47). However, more studies are needed to establish these findings. Alternatively, an indirect effect resulting from vaccine-induced trained immunity could contribute to bystander protection. In fact, studies have reported protection against SARS-CoV-2 using BCG vaccination (48, 49). It is possible that ^Flu^VLP might have acted as an adjuvant before infection resulting in a better bystander effect upon challenge with the SARS-CoV-2. We have recently shown that ^Flu^VLP induces a cross-protective effect against heterologous influenza viruses without inducing hemagglutination inhibiting antibody response (50). Therefore, further studies are needed to understand the potential cross-protection between influenza and SARS-CoV-2 and the associated mechanisms.

Our study has some limitations. Despite demonstrating an effective protection by ^Flu^VLP-RBD hybrid vaccine in rhesus macaques there are potential caveats in this study. First the number of animals in each group was statistically less based on the power analysis. Therefore, statistical significance was not reached despite effective protection in ^Flu^VLP-RBD vaccinated macaques. Secondly, there was no concurrent control group in this study. Though the historical ancestral control group used in the current study was challenged with the same SARS-CoV-2 strain (WA1/2020) and the route of viral challenge (nasal and tracheal), a higher dose (10^5^ pfu) and different viral stocks were used in this study. Nevertheless, the current study shows that an influenza-based SARS-CoV-2 hybrid vaccine induces robust antibody responses against influenza and multiple SARS-CoV-2 variants, efficiently protects rhesus macaques against SARS-CoV-2 (WA1/2020) and prevents severe lung pathology. These findings provide key insight for SARS-CoV-2 and influenza dual vaccination strategy for human clinical trials.

## Material and Methods

### Construction and characterization of influenza-VLP based dual vaccine

Vaccines were constructed and characterized as previously described (31). We used human GM-CSF (instead of monkey GM-CSF) for developing the fusion protein with RBD in the VLP hybrid vaccine since human GM-CSF works in nonhuman primates. Similarly, human IL-12 was used in the vaccine. Influenza A/Puerto Rico/8/1934 (PR8) strain (^Flu^VLP) expressing codon-optimized hemagglutinin (H1 HA) and matrix M1 proteins was produced in Sf9 insect cells and purified by tangential flow diafiltration and anion exchange (Capto Q) chromatography by Medigen (Frederick, MD, USA). GPI-RBD-hGM-CSF recombinant fusion protein and GPI-hIL-12 were affinity purified from CHO-S cells (Invitrogen) stably expressing the proteins using anti-human GM-CSF (clone BVD2-21C11) BioXCell (Lebanon, NH, USA)) and anti-human IL-12 (Clone C8.6, produced in-house at Metaclipse Therapeutics) antibodies coupled with NHS-Sepharose affinity columns. The purity and identity of the fusion protein was confirmed by colloidal blue and western blot. Purified GPI-RBD-hGM-CSF and hGPI-IL12 were incorporated into the ^Flu^VLP by incubating it with the GPI-anchored fusion proteins for 2 h at 37°C. A total of 1.0 mg of ^Flu^VLP was incubated with 200 μg of purified GPI-RBD-hGM-CSF and 25 μg of purified GPI-hIL-12. Unincorporated GPI-proteins were washed off by ultracentrifugation at 210,000g at 4°C for 1 hr. The resulting ^Flu^VLP-RBD vaccine pellet was resuspended in DPBS and characterized for incorporation of GPI-proteins by flowcytometry using anti-hGM-CSF-APC (Clone BVD2-21C11, Biolegend, Catalog# 502310) and anti-hIL-12-PE (Cat# 554575, BD Biosciences) antibodies. The ACE2-binding property of fusion RBD was determined by ELISA. Briefly, ELISA plate was coated with GPI-hGM-CSF-RBD fusion protein, blocked with PBS with 3%BSA and then incubated with various concentrations of biotinylated ACE-2 which was detected by adding streptavidin-HRP and TMB substrate as described in our earlier studies (Bommireddy et al 2022).

### Immunization of rhesus macaques

Young (3-4 years old) adult Indian rhesus macaques (RMs) from the Emory Primate Center’s breeding colony were randomly selected as shown table S1 following the Emory University’s Institutional Animal Care and Use Committee approval and guidelines. All animals were treated in accordance with the US Department of Agriculture regulations and recommendations derived from the Guide for the Care and Use of Laboratory Animals. Eight RMs were equally divided into two treatment groups. Group 1 received influenza A/PR8 virus like particles (^Flu^VLP) and group 2 received ^Flu^VLP-RBD vaccinations. A total of 25μg of ^Flu^VLP and ^Flu^VLP-RBD were subcutaneously delivered at week 0 (prime) and week 4 (boost) and immune response was followed in blood at week 0, 2, and 4 of each vaccination. RMs were challenged with 1 x 10^5^ pfu of SARS-CoV-2 WA-1/2020 strain via intranasal and intratracheal delivery of challenge stock at week 8 as described before (32, 33). Viral infection was followed in lungs (using bronchoalveolar lavage), nasopharynx, and throat at day 0, 2, 4, 7, and 10 of infection for preclinical assessment of protection.

### SARS-CoV-2 WA-1/2020 infection quantification

Viral infection was estimated by quantifying SARS-CoV-2 sub-genomic RNA in bronchoalveolar lavage (BAL), nasopharyngeal (NP) swabs, throat swabs at day 0, 2, 4, 7, and 10 of challenge as described before(32, 33). Briefly, swabs were collected in 1mL of Viral Transport Medium (VTM; Labscoop (VR2019-1L)) and viral RNA was extracted using the QiaAmp Viral RNA mini kit according to the manufacturer’s protocol. Quantitative PCR (qPCR) was performed using the primer and probe sequences for the mRNA transcripts of the E gene (51). The primer and probe sequences used were SGMRNA-E-F: 5′-CGATCTCTTGTAGATCTGTTCTC-3′, SGMRNA-E-R: 5′-ATATTGCAGCAGTACGCACACA-3′, and SGMRNA-E-Pr: 5′-FAM-ACACTAGCCATCCTTACTGCGCTTCG-3′. qPCR reactions were performed in duplicate with the Thermo-Fisher 1-Step Fast virus mastermix using the manufacturer’s cycling conditions, 200nM of each primer, and 125nM of the probe. The limit of detection in this assay was about 128 copies per mL of VTM/BAL depending on the volume of extracted RNA available for each assay. To verify sample quality the CDC RNase P p30 subunit qPCR was modified to account for rhesus macaque specific polymorphisms. The primer and probe sequences are RM-RPP30-F 5′-AGACTTGGACGTGCGAGCG-3′, RM-RPP30-R 5′-GAGCCGCTGTCTCCACAAGT-3′, and RPP30-Pr 5′-FAM-TTCTGACCTGAAGGCTCTGCGCG-BHQ1-3′ (52). A single well from each extraction was run as described above to verify RNA integrity and sample quality via detectable and consistent cycle threshold values (Ct between 25-32).

### Binding antibody responses using ELISA

Binding IgG against RBD of SARS-CoV-2 WA-1/2020, Delta, and Omicron variants or influenza-VLP in serum was quantified by enzyme-linked immunosorbent assay (ELISA) as described previously (53). Briefly, ELISA plates (Nunc) were coated with 2mg/ml of recombinant RBD proteins or 200ng/well influenza A/PR8-VLP in Dulbecco’s phosphate-buffered saline (DPBS) and incubated overnight at 4°C. Next day, protein coated plates were blocked with 5% milk and 4% whey powders in DPBS supplemented with 0.05% Tween-20 for 2h at room temperature (RT). Blocking buffer was discarded and serum samples were added to the wells with increasing dilution (starting dilution was 1 in 100, subsequent 3-fold serially diluted). Plates were incubated for 2h at RT followed by washing. Total RBD-binding IgG was detected using HRP-conjugated anti-monkey IgG secondary antibody (dilution factor - 1:10, 000), for 1h at RT. Finally, ELISA plates were washed, and chromogen was developed using TMB (2-Component Microwell Peroxidase Substrate Kit), which was followed by stopping the reaction with the addition of 1N phosphoric acid solution. Plates were read at 450 nm wavelength using the plate reader (Molecular Devices; San Jose, CA, USA). ELISA endpoint titers were defined as the highest reciprocal serum dilution that yielded an absorbance > 2-fold over background values.

### Live-virus neutralization

icSARS-CoV-2 (closely resembling the original Wuhan strain), B.1.617.1 and B.1.1.529 variants were plaque purified and propagated once on VeroE6-TMPRSS2 cells. All viruses used in this study were deep sequenced and confirmed as described (54, 55).

Vero-TMPRSS2 cells were cultured in complete DMEM medium consisting of 1x DMEM (VWR, #45000-304), 10% FBS, 2mM L-glutamine, and 1x antibiotic as previously described (54, 56). Samples were diluted at 3-fold in 8 serial dilutions using DMEM in duplicates with an initial dilution of 1:10 in a total volume of 60 ul. Serially diluted samples were incubated with an equal volume of SARS-CoV-2 (100-200 foci per well) at 37C for 1 hour in a round bottomed 96-well culture plate. The antibody-virus mixture was then added to Vero cells and incubated at 37C for 1 hour. Post-incubation, the antibody-virus mixture was removed and 100 μl of prewarmed 0.85% methylcellulose overlay was added to each well. Plates were incubated at 37C for 18 to 40 hours, and the methylcellulose overlay was removed and washed six times with PBS. Cells were fixed with 2% paraformaldehyde in PBS for 30 minutes. Following fixation, plates were washed twice with PBS, and permeabilization buffer (0.1% BSA, 0.1% Saponin in PBS) was added to permeabilize cells for at least 20 minutes. Cells were incubated with an anti-SARS-CoV spike primary antibody directly conjugated to Alexa Fluor-647 (CR3022-AF647) overnight at 4°C. Cells were then washed twice with 1x PBS and imaged on an ELISPOT reader (CTL Analyzer). Antibody neutralization was quantified by counting the number of foci for each sample using the Viridot program (57). The neutralization titers were calculated as follows: 1 – (ratio of the mean number of foci in the presence of sera and foci at the highest dilution of the respective sera sample). Each specimen was tested in duplicate. The FRNT-50 titers were interpolated using a 4-parameter nonlinear regression in GraphPad Prism 9.2.0. Samples that do not neutralize at the limit of detection at 50% are plotted at 20 and used for geometric mean and fold-change calculations.

### Antibodies

Anti-human antibodies known to cross-react with rhesus macaques were used. For T cell phenotype and cytokine assay, anti-CD3 (Clone SP-34-2; BD Biosciences), anti-CD8 (SK1; BD Bioscience), anti-CD4 (L200; BD Biosciences), anti-IFNγ (B27, BD Biosciences), anti-TNFα (MAb11, BD Biosciences), and anti-IL-2 (MQ1-17H12, BD Biosciences) antibodies were used. MM57 is an RBD-specific mouse monoclonal antibody shown to neutralize pseudovirus infection in vitro. Anti-RBD mAb (clone MM57) was obtained from Sino Biologicals (Cat#40592). Streptavidin-HRP was obtained from R&D Systems. Biotinylated human ACE2 from ACRO Biosystems (Cat#AC2-H82F9), Peroxidase (HRP)-conjugated goat anti-mouse IgG F(ab’)2 specific antibody was from ThermoFisher Scientific/Pierce (Cat#31436). FITC-conjugated streptavidin was purchased from BD Biosciences (Cat#554060).

### Intracellular cytokine staining (ICCS) assay and flowcytometry

Antigen specific CD4 and CD8 T cells were analyzed by measuring cytokine responses against SARS-CoV-2 spike (S) peptides as described before (32, 33). Briefly, peripheral blood mononuclear cells (PBMCs) at week 0, 2 of vaccinations and day 10 of infection were stimulated using overlapping S-peptide pools (BEI resources; NR-52402). 2 x 10^6^ PBMCs were stimulated with 1 μg/ml peptide pool in complete RPMI containing 1μg/ml each of CD28 (Cat#555725) and CD49d (Cat#555501) co-stimulatory antibodies. In addition, 2 x 10^6^ PBMCs were also stimulated each with DMSO (negative control) and PMA (80ng/ml) plus ionomycin (1μg/mL). Cells were incubated at 37°C for 2 hours(h) prior to addition of Golgistop/Golgiplug followed by additional incubation at 37°C for 4h. Cells were stained to measure T cell cytokine response to S. Cells were washed with FACS wash (1X PBS, 2% FBS and 0.05% sodium azide) followed by surface staining with Live/Dead-APC-Cy7, CD3-PerCP-Cy5.5, CD4-BV650 and CD8-BUV496. Cells were incubated for 20 minutes at 4°C followed by FACS wash. Washed cells were permeabilized and fixed with Cytofix/Cytoperm for 15 minutes at 4°C followed by perm-wash. Permeabilized cells were stained with IFN γ, TNFα, and IL-2 and incubated for 20 minutes at 4°C followed by two perm-wash and one FACS wash. Flow cytometry was performed using BD LSR Fortessa flow cytometer and data was analyzed using FlowJo software.

### Histopathological examination

Lung pathology was assessed as previously described (32, 33). Briefly, lung lobes were harvested at necropsy and fixed with 10% neutral-buffered formalin for 24 hours at room temperature, paraffin-embedded, sectioned at 4 μm, and stained with H&E. The H&E slides were examined by a board-certified veterinary pathologist in a blinded fashion. For each animal, all the lung lobes were used for analysis, and affected microscopic fields were scored semi quantitatively as grade 0-5 (mild), 5-10 (moderate), >10 (severe). Scoring was performed on the basis of these criteria: number of lung lobes affected, type 2 pneumocyte hyperplasia, alveolar septal thickening, fibrosis, perivascular cuffing, peribronchiolar hyperplasia, inflammatory infiltrates, and hyaline membrane formation. Total lung pathology was calculated by combining scores from each criterion.

### Quantification and statistical analysis

Statistical significance was calculated using a two-tailed unpaired nonparametric Mann-Whitney test. Correlations were analyzed using Spearman r method. GraphPad Prism version 8.4.3 (GraphPad Software) was used to perform data analysis and statistical tests.

## ACKNOWLEDGMENTS

We thank the Division of Pathology and Animal Resources at ENPRC for their help in ABSL3 during the virus challenge. BioRender was used to make the schematic shown in Figure 2A. The content is solely the responsibility of the authors and does not necessarily reflect the official views of the National Institute of Health. The authors thank Allison N. Blackerby, Dominique N. Gales-Badea, Swe-Htet Naing, and Lahcen Jaafar for GPI-cytokine purification. We thank Dr. Kristen M. Jacobsen for critical reading and editing the manuscript. We also thank Brenda Wehrle, Zeba Liyakat Momin, Kheyanna Davis and Traci H. Legere for help with sample processing, shipping, and ordering reagents.

## AUTHOR CONTRIBUTIONS

Conceptualization: RRA, PS, CDP, RB, SAR, SR, and MSS; Methodology: SAR, RB, NKR, LL, CDP, SR, MSS, SR, PS, and RRR; Investigation: SAR, RRA, RB, NKR, LL; Funding acquisition: RRA, PS, SJCR, and CDP; Project administration: RRA, CDP, SR, and PS; Supervision: RRA; Writing—original draft: SAR, and RRA and PS; All authors have read and agreed to the published version of the manuscript.

